# Evaluation of the antimicrobial, anti-adhesion, anti-biofilm and cell proliferation assay of a de-novo goji berry extract against periodontal pathogens: a comparative in-vitro study

**DOI:** 10.1101/2022.03.07.483227

**Authors:** Amee Dilip Sanghavi, Aditi Chopra, Ashmeet Shah, Richard Lobo, Padmaja A Shenoy

**Affiliations:** Department of Periodontology, Manipal College of Dental Sciences, Manipal Academy of Higher Education, Manipal, Karnataka, India, Pin: 576104; Department of Pharmacognosy, Manipal College of Pharmaceutical Sciences, Manipal Academy of Higher Education, Manipal, Karnataka, India, Pin: 576104; Department of Microbiology, Kasturba Medical College, Manipal Academy of Higher Education, Manipal, Karnataka, India, Pin: 576104

**Keywords:** Goji Berry, Lycium barbarum, Periodontitis, Antimicrobial, Herb, MTT assay

## Abstract

*Lycium barbarum*, commonly known as goji berry, Chinese berry, or Tibetian berry, is emerging as a popular “superfood” with anti-inflammatory and antioxidant properties. Goji berry is being used for the treatment of various cancers, gastrointestional disorders, stroke, diabetes, Alzheimer’s disease, and glaucoma. However, its use for management of oral inflammatory diseases has not been explored. Therefore, the present study aims to evaluate the antimicrobial, anti-adhesion, and anti-biofilm, and cytotoxic properties of an ethanolic extract of *Lycium barbarum* (LBE) against oral and periodontal pathogens. The antimicrobial properties of LBE against five microorganisms were tested and compared against Chlorhexidine and doxycycline along with cytotoxicity and cell viability on the gingival fibroblast and modified keratinocyte cell lines. The anti-adhesion and anti-biofilm properties of LBE against *Porphyromonas gingivalis*, at its minimal bactericidal value, were evaluated. The antimicrobial, anti-adhesion and antibiofilm properties of LBE were found to be comparable to chlorhexidine but less than that of doxycycline. The LBE extract was also compactible to gingival fibroblast tissues and oral keratinocytes at 1 mg/ml. The results proved that goji berry is as effective as chlorhexidine and can be used as a promising natural herb for the management of inflammatory diseases of oral cavity.

## 1. Introduction

Periodontitis is a chronic inflammatory disease caused by the interaction of pathogenic microorganisms in the oral microbiome to soft tissues surrounding the tooth (Chapple et al., 2018). This host-microbial interaction in the periodontal tissues causes a release of various pro-inflammatory cytokines, microbial by-products, and free radicals in the oral tissues. These chemical mediators and microbial by-products initiate an inflammatory response that causes gingivitis. If gingivitis is not controlled, the inflammation spreads to the underlying tissues and result in the onset of periodontitis. Chronic periodontitis results in bone loss, pocket formation, tooth mobility, with partial or complete loss of teeth (Chapple et al., 2018). The periodontal microorganisms can enter the systemic circulation and increase the risk of Diabetes mellitus, atherosclerosis, chronic kidney disease, Alzheimer’s, poor pregnancy outcomes, rheumatoid arthritis, and oropharyngeal and pancreatic cancer (Winning and Linden, 2015). Since the global burden of periodontal disease is rapidly increasing, it is important to develop effective therapeutic modalities to control inevitably periodontal inflammation and its associated systemic complications (Nazir et al., 2020).

The most common treatment modality to control periodontal diseases is the mechanical debridement of oral biofilm and calculus from the surface of the teeth with or without antimicrobial agents (Smiley et al., 2015). Antibiotics are the most common adjunct used along with scaling and root planing for the management of gingivitis and periodontitis (Bonito et al., 2006; Zandbergen et al., 2013; Keestra et al., 2015). However, with the wide-spread use of antibiotics, many oral pathogens have now become resistant to commonly prescribed antibiotics (Bell et al., 2014). Therefore, there is a changing trend of using herbal or plant-based products for the management of gingival and periodontal diseases (Karygianni et al., 2015).

Plant-based products have shown to possess good anti-inflammatory, astringent, antioxidant, and antimicrobial properties against oral pathogens. Green tea, curcumin, cranberry, tulsi, guava, aloe vera, neem, pomegranate, mango leaf. have shown good antimicrobial properties against the oral and periodontal pathogens (Bautista-Pérez et al, 2004; Kushiyama et al, 2009; La et al, 2010; Bhat et al, 2011; Kumar et al, 2013). The extracts from these herbs have been incorporated into gels, mouthwash, tablets, powder, local drug delivery agents for the treatment of gingivitis and periodontitis.

Recently, *Lycium barbarum*, commonly known as goji berry, Chinese berry, or Tibetian berry, or wolfberry, has emerged as a potential antioxidant and antimicrobial agent for the management of inflammatory conditions. *Lycium barbarum* is a tree that belongs to the family of Solanaceae. It is native to southeast Europe and Asia. *Lycium barbarum* is regarded as a “superfood” as it contains various beneficial phytochemical elements. Bondia-Pons et al. (2014) and Forino et al. (2016) stated that the fruits of *Lycium barbarum* contain important constituents like *Lycium barbarum* Polysaccharides or LBPs, carotenoid (zeaxanthin dipalmitate), vitamins, and flavonoids (catechin, epicatechin, quercetin, quercetin-rhamno-di-hexoside, and quercetin-3-O-rutinoside aglycones myricetin) and kaempferol p-coumaric acid, rutin, caffeic acid, scopoletin, N-trans-feruloyl tyramine, Ncis-feruloyl tyramine, N-feruloyl tyramine dimer, linoleic acid and, kaempferol glycosides, isomers of dicaffeoylquinic acid and coumaric acid (Amagase & Farnsworth (2011). Donno et al. (2015). It also contains phenolic acid (chlorogenic acid), monoterpenes (phellandrene, sabinene, γ-terpinene), organic acids (citric acid, malic acid, oxalic acid, quinic acid, tartaric acid), and vitamin C analogs. Various preclinical and clinical studies conducted by Toyoda-Ono et al, 2004; Chen et al, 2009; Low et al, 2019; Xiao et al, 2019 have proven the medicinal, therapeutic, and health-promoting properties of *Lycium barbarum*. Lycium barbarum has been tried for the treatment of aging, fatigue, cancer, colitis, stroke, diabetes, Alzheimer’s disease, and glaucoma (Mocan et al., 2017, Skenderidis et al., 2019; Pires et. al, 2018). However, its effect of the oral microorganisms has not been explored. Therefore, the present study aims to investigate the antimicrobial, anti-adhesion, and anti-biofilm, and cytotoxic properties of ethanolic extract of *Lycium barbarum* fruit against oral and periodontal pathogens. The results of the paper are of paramount importance as it proves for the first time the efficacy of Lycium barbarum against oral and periodontal pathogens and provide a preliminary data the antimicrobial properties of Goji berry extract. Since the global burden of periodontal disease is rapidly increasing and many oral bacteria are becoming resistance to antibiotics, it is important to develop natural therapeutic modalities to control inevitably periodontal inflammation and its associated systemic complications.

## 2. Experimental methodology

### 2.1 Preparation of the ethanolic extract

The fruits of Goji berry (Lycium barbarum. L; family Solanaceae) were procured from an authorized distributor (Kenny delights Pvt. limited) which were imported from China. The fruits were identified and authenticated by Dr. Gopalkrishna Bhat, a botanist and taxonomist and a retired Professor from Poornaprajna College, Udupi. The voucher specimen bearing a reference number [PP626] was deposited in the Department of Pharmacognosy, Manipal College of Pharmaceutical Sciences, Manipal Academy of Higher Education (MAHE), Manipal. The research was done in Compliance with Ethical Standards after obtaning the Institutional Ethics committee approval, KMC Manipal (IEC: 884/2018)

The fruit was washed with distilled water and dried in a hot air oven at a temperature of 45°C. The dried berries were coarsely powdered in a mixture grinder, and about 250g of the powdered berries were obtained. The powder was then macerated with 1000ml of ethanol for 3 days with occasional shaking. The macerated powder was subsequently filtered, and the solvent was removed with a rota evaporator. The extract was carefully collected and stored in a desiccator until further investigations and analysis. Upon desiccation, an extract with a semisolid sticky, brown color mass with a sweet fruity fragrance was obtained.

### 2.2 Evaluation of the antimicrobial properties

The bacterial cultures from the repository of Dr. Kishore G. Bhat Maratha Mandal’s NGH Institute of Dental Sciences and Research Centre, Belgaum, Karnataka, India were used. The bacterial strains used for the present study were as follows: Streptococcus mutans (Sm): ATCC 10449; *Porphyromonas gingivalis* (Pg): ATCC 33277; *Aggregatibacter actinomycetemcomitans* (Aa): ATCC 700685; *Fusobacterium nucleatum* (Fn): ATCC 23726; *Prevotella intermedia* (Pi): ATCC 25611; *Tanerella forsythia* (Tf): ATCC 43037. The antimicrobial properties were checked by the following test:

#### 2.2.1. Minimum Inhibitory Concentration (MIC) (Figure 1 and Table 1)

**Figure 1:**
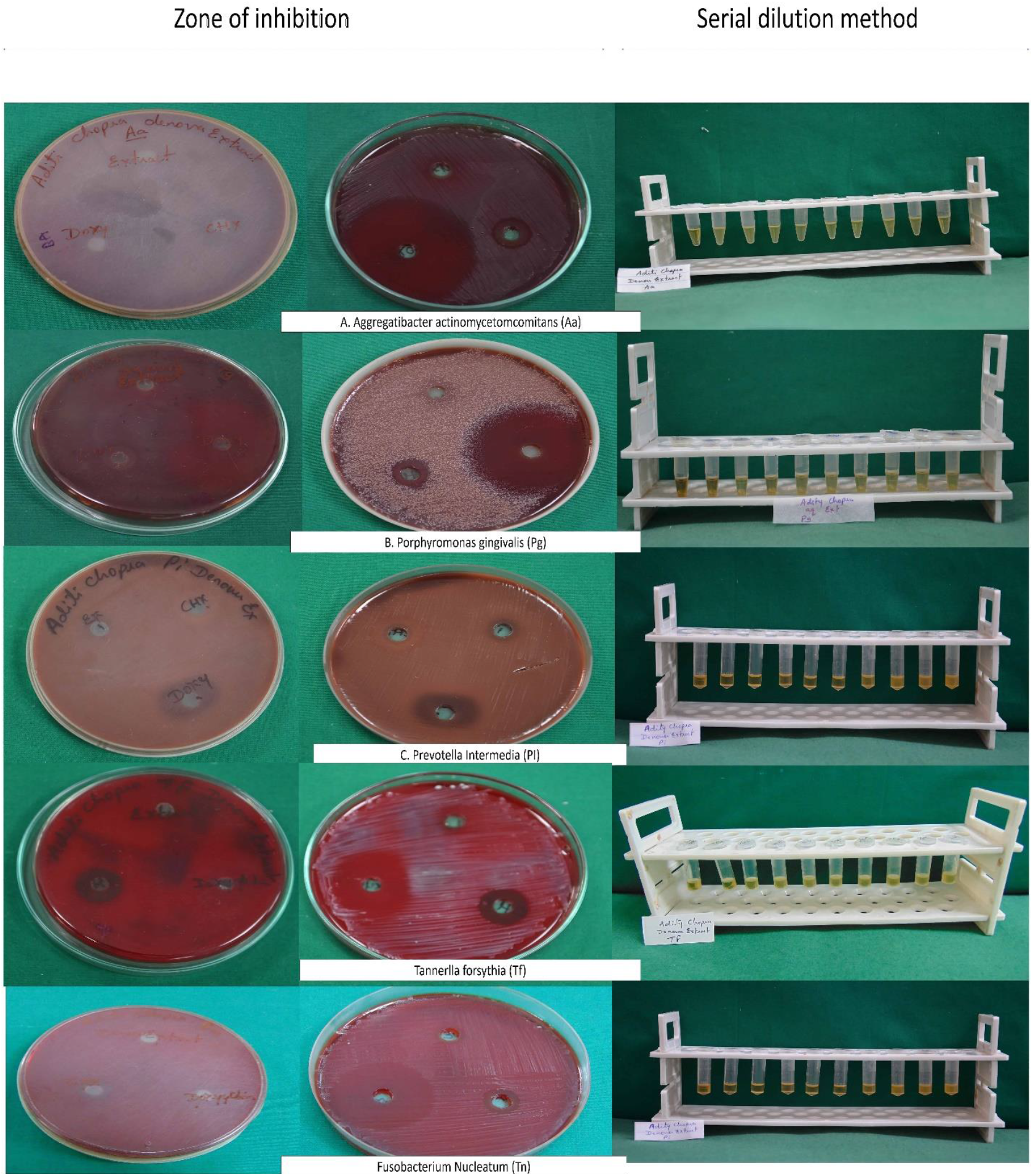
Mean zones of inhibition by Lycium barbarum extract compared to doxycycline and chlorhexidine against Aggregatibacter actinomycetemcomitans (Aa), Porphyromonas gingivalis (Pg), Prevotella intermedia(Pi), Tannerlla forsythias (Tf), Fusobacterium nucleatum (Fn)

**Table 1:**
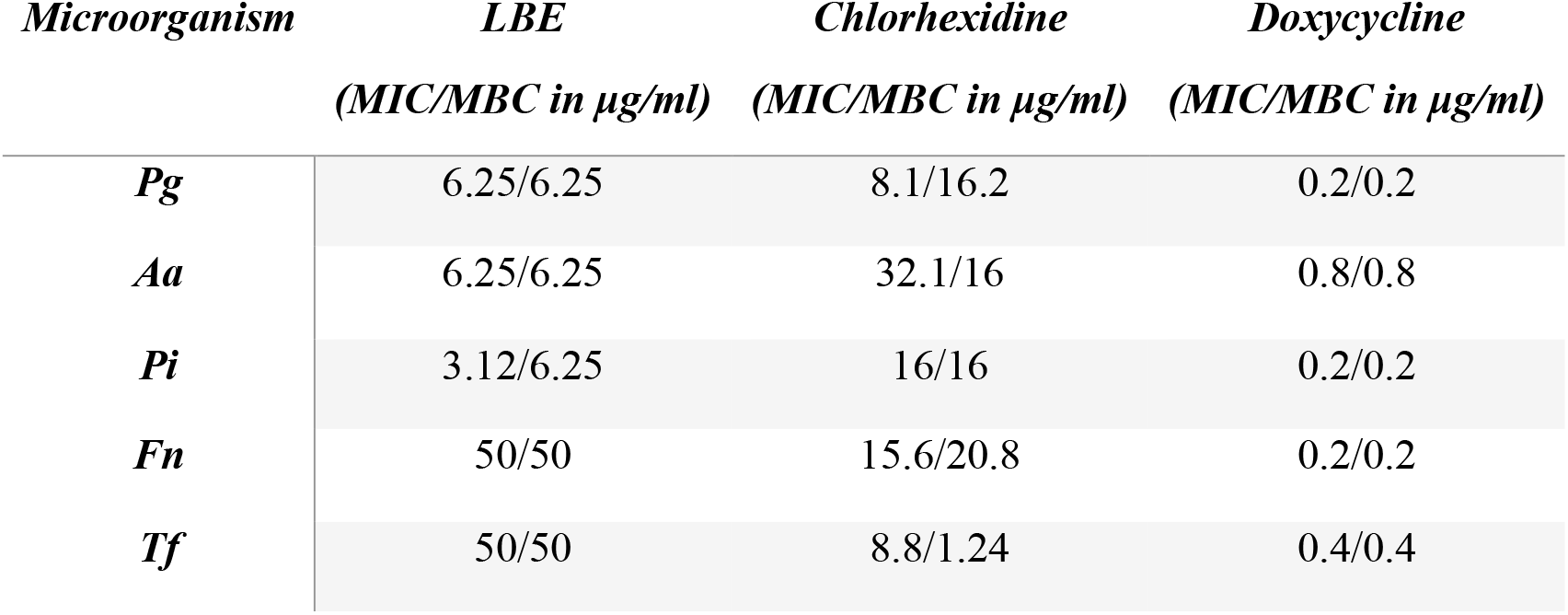
Minimal Inhibitory concentration (MIC)/ Minimal bactericidal concentration (MBC) of the ethanolic extract of Lycium barbarum compared to chlorhexidine and doxycycline.

| *Microorganism* | *LBE*<br>(MIC/MBC in $\mu\text{g/ml}$) | *Chlorhexidine*<br>(MIC/MBC in $\mu\text{g/ml}$) | *Doxycycline*<br>(MIC/MBC in $\mu\text{g/ml}$) |
| --- | --- | --- | --- |
| *Pg* | 6.25/6.25 | 8.1/16.2 | 0.2/0.2 |
| *Aa* | 6.25/6.25 | 32.1/16 | 0.8/0.8 |
| *Pi* | 3.12/6.25 | 16/16 | 0.2/0.2 |
| *Fn* | 50/50 | 15.6/20.8 | 0.2/0.2 |
| *Tf* | 50/50 | 8.8/1.24 | 0.4/0.4 |

The MIC was determined by using the ‘microdilution technique’ with thioglycollate broth (Himedia Pvt. Ltd., Mumbai, India) (ref). In the initial tube, 20 μl of the extract was added into the 380 μl of thioglycollate broth. For dilutions, 200 μl of thioglycollate broth was added into the next 9 tubes separately. From the initial tube, 200 microliters were transferred to the first tube containing 200 microliters of thioglycollate broth. This was considered a 10-1 dilution. From a 10-1 diluted tube, 200 μl were transferred to the second tube to make 10-2 dilution. Similarly, the serial dilution was repeated up to 10-9 dilution. From the maintained stock cultures of the microorganisms (Sm, Pg, Aa, Fn, Pi, Tf), a microliter was taken from each and added into 2 ml of thioglycollate broth. In each serially diluted tube, 200 μl of above-cultured suspension was added. The tubes were then incubated for 48-72 hours in an anaerobic jar at 37°C and observed for turbidity. The turbidity was checked using optical density and the reading was observed at 450 nm. The experiment was done in triplicate for each strain.

#### 2.2.2. Minimum Bactericidal Concentration (MBC)

From the MIC dilutions tubes, the first 5 tubes were plated (which was sensitive in MIC) and incubated for 24 hours. The next day, the total colony count was taken. The MBC was conducted to evaluate whether there was a bacteriostatic or bactericidal effect of the LBE against the periodontal pathogens. The lowest concentration of the subculture with no growth was considered the minimum bactericidal concentration. The experiment was done in triplicate for each strain (Table 1).

#### 2.2.3 Well- Diffusion Assay (Table 2 and Figure 1)

**Table 2:**
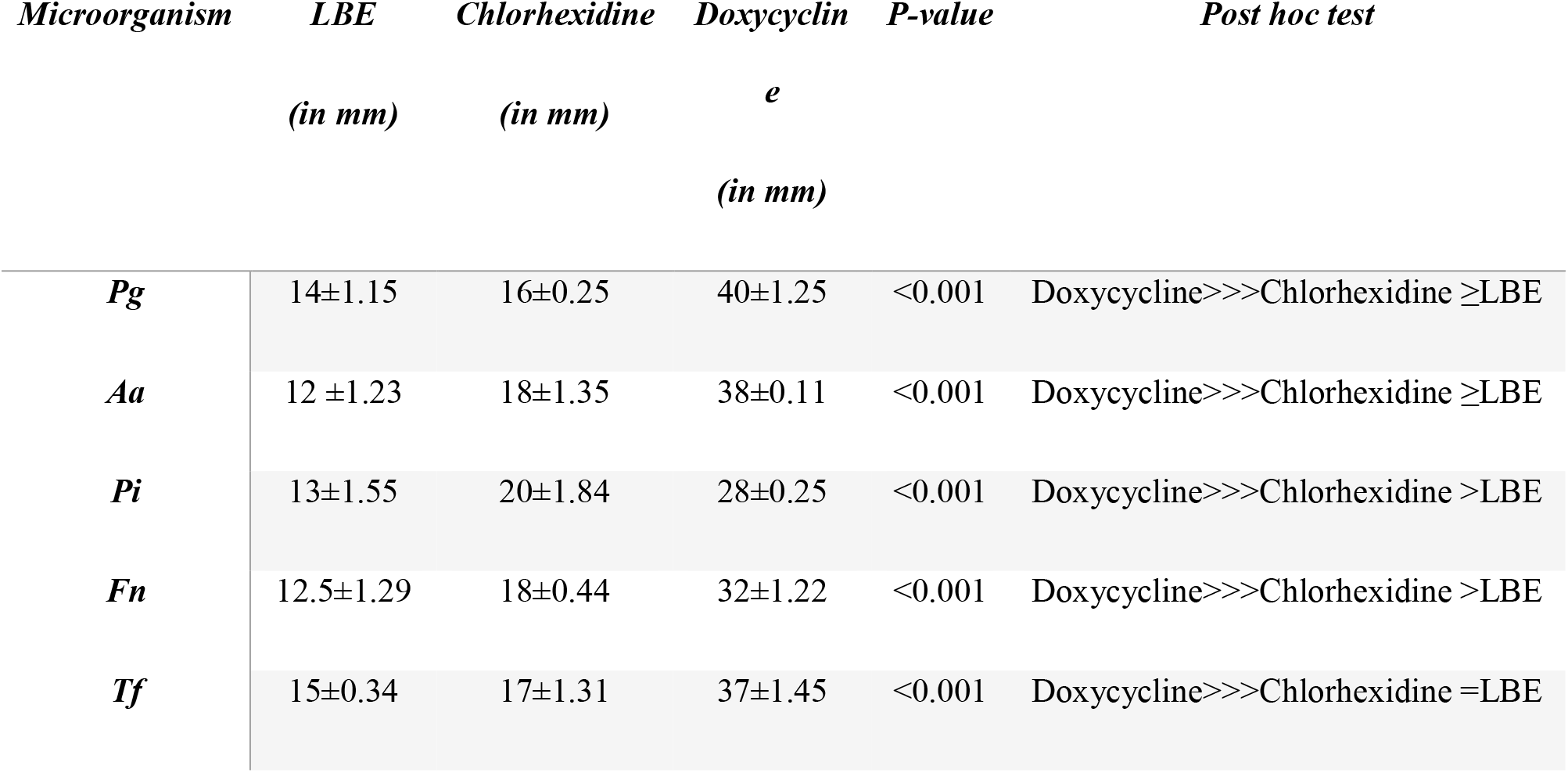
**Mean zones of inhibition by Lycium barbarum extract compared to doxycycline and chlorhexidine against Aggregatibacter actinomycetemcomitans (Aa), Porphyromonas gingivalis (Pg), Prevotella intermedia(Pi), Tannerlla forsythias (Tf), Fusobacterium nucleatum (Fn) [One-way variance of ANOVA and post-hoc-Tukey)**

| *Microorganism* | *LBE*<br>(in mm) | *Chlorhexidine*<br>(in mm) | *Doxycyclin*<br>*e*<br>(in mm) | *P-value* | *Post hoc test* |
| --- | --- | --- | --- | --- | --- |
| *Pg* | 14±1.15 | 16±0.25 | 40±1.25 | <0.001 | Doxycycline>>>Chlorhexidine ≥LBE |
| *Aa* | 12 ±1.23 | 18±1.35 | 38±0.11 | <0.001 | Doxycycline>>>Chlorhexidine ≥LBE |
| *Pi* | 13±1.55 | 20±1.84 | 28±0.25 | <0.001 | Doxycycline>>>Chlorhexidine >LBE |
| *Fn* | 12.5±1.29 | 18±0.44 | 32±1.22 | <0.001 | Doxycycline>>>Chlorhexidine >LBE |
| *Tf* | 15±0.34 | 17±1.31 | 37±1.45 | <0.001 | Doxycycline>>>Chlorhexidine =LBE |

Agar well diffusion method using Brain Heart Infusion agar (Himedia Pvt. Ltd., Mumbai, India) was used to test the antimicrobial activity of the extract against the pathogens. The agar plates were brought down to room temperature after which, the colonies of microorganisms were inoculated on the agar plates using a swab. The facultative anaerobes were incubated in tubes at 37°C for 48-72 hours in a carbon dioxide jar. The strict anaerobes were incubated in anaerobic jars for 48-72 hours before the experimentation. Within 15 min, the inoculum was adjusted to a McFarland 0.5 turbidity standard and the suspension was regulated with a photometric device. After the process of inoculation, a sterile cotton swab was dipped into the inoculum and rotated against the wall of the tube above the liquid to remove the excess inoculum. The entire surface of the agar plate was swabbed three times and the inoculum was transferred. The plates were subsequently rotated by approximately 60° between streaks to ensure even distribution. The inoculated culture plates were allowed to stand for at least 3 minutes but not more than15 minutes. The procedure of inoculum preparation and inoculation of culture media was done for Sm, Pg, Aa, Fn, Tf, and Pi on five agar plates for five respective concentrations (0.5%, 1%, 2%, 5%, and 10%) of the LBE. The stock solution was prepared by using 10mg of the compound and dissolved in 1 ml of Dimethyl Sulfoxide (DMSO). A hollow tube of 5mm diameter was heated and then pressed on the inoculated agar plate and then removed immediately. This results in good formation on the agar plates and a total of 5 wells were made on each plate. A micropipette was used to add 75μl, 50μl, 25μl, 10μl, and 5μl in each well. The plates were then incubated for a period of 24 h at 37° C. After the incubation period, plates were ready only if the lawn of growth was confluent or nearly confluent. The diameter of the inhibition zone was measured to the nearest whole millimeter by using a Vernier Caliper. The microbiological procedure was repeated 4 times for each bacterium, and the corresponding four values of zones of inhibition for each concentration of *Lycium barbarum* extract were obtained. The zone of inhibition was compared to doxycycline, chlorhexidine and control.

### 2.3. Cell proliferation assay (MTT assay) for evaluating the cytotoxicity and cell viability on using gingival fibroblast and keratinocyte cell line (Figure 2)

**Figure 2:**
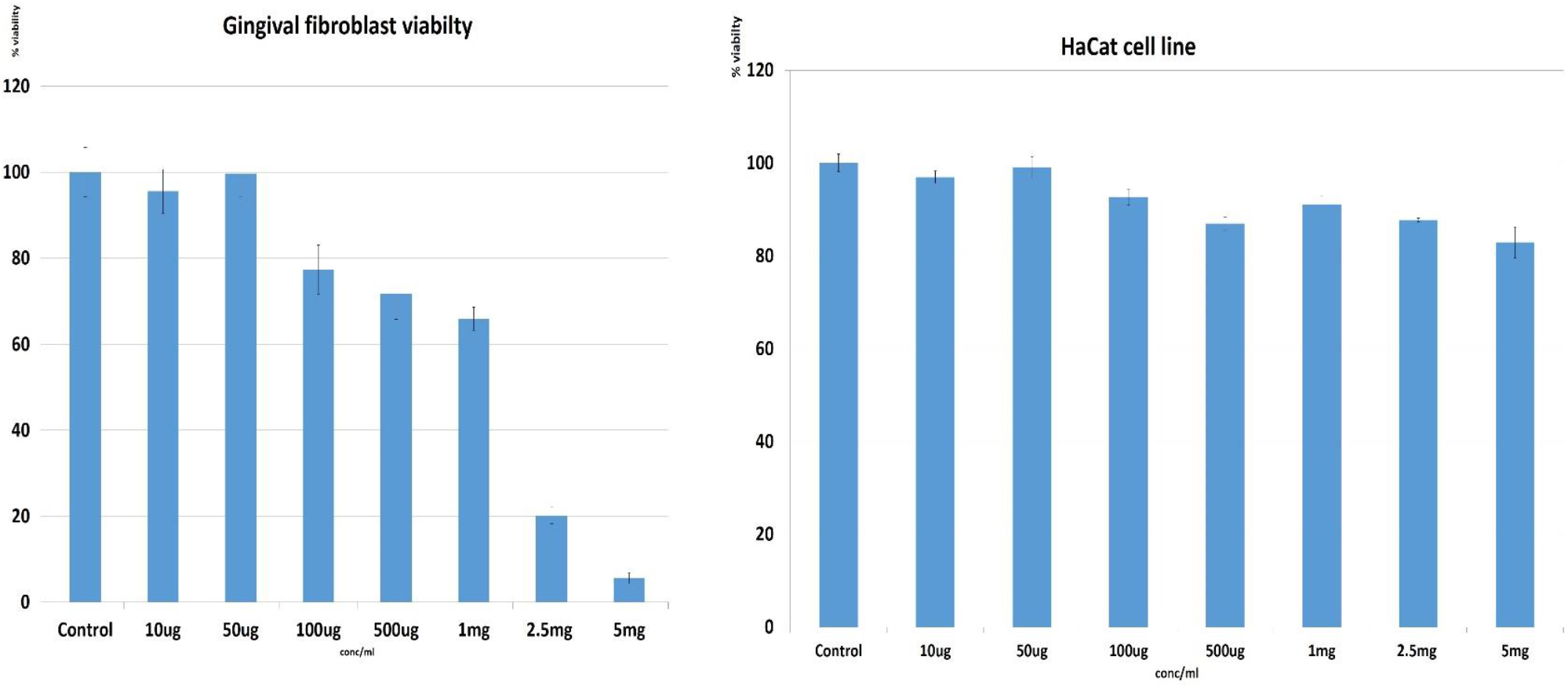
Cell proliferation assay (MTT assay) of LBE against the gingival fibroblasts and modified Keratinocye cell line (HACAT): The results showed that LBE maintained the cell viability of more than 75 % of the gingival fibroblast and 80% of modified keratinocyte cell line (HCAT) at a concentration of 1mg, and 5 mg respectively compared to control (normal saline)

The cell viability was measured using gingival fibroblast (HGF-1) and keratinocyte cell line (HACAT transformed keratinocyte) using the MTT assay. The human-derived fibroblast and keratinocyte cell line cells were exposed to the extract at different concentrations for 48 hours. The dose-dependent inhibition of the *Lycium barbarum* on cell proliferation was observed. The gingival fibroblast/ HACAT transformed keratinocyte was seeded at 1 × 10^5^ cells/mL in 96 well microtiter plates in Minimum Essential Medium with fetal bovine serum. The cells were incubated overnight for attachment. The drug concentrations in serial three-fold dilutions were added in triplicates and incubated for 48h at 5% CO2 at 37°C. The cells were treated with the extract at varying concentrations 10ug, 50ug, 100ug, 500ug, 1mg, 2.5mg, 5mg and compared to 5 µg of positive control (normal saline) for 48 hours, respectively. MTT dye (5 mg/mL) was added to each well for at least 4 hours of treatment. Thereafter, the cells were treated with 3-[4,5-dimethylthiazol-2-yl]-2,5-diphenyltratrazolium bromide (MTT) (Sigma Chemical Co., St. Louis, MO). Four hours later, all of the medium including MTT solution (5 mg/mL) was aspirated from the wells. The remaining formazan crystals were dissolved in DMSO and the absorbance was measured at 570 nm using a 96 well microplate reader (Synergy TM HT, Bio-Tek Instruments, Inc). The cytotoxicity index was determined using the untreated cells. The percentage of cytotoxicity was calculated using the background-corrected absorbance as follows: % cytotoxicity = (1−absorbance of the experimental well) absorbance of negative control well×100).

### 2.4 Anti-adhesion Assay

The anti-adhesion activity of the extract was tested against Pg, at its MBC value. A 96-well flat bottomed microtitre plate was taken and wells were loaded with 100 μl of test samples (Goji, CHX, & amp; blank media) in triplicates. Pg culture (100 μl) was added to each well so that a total volume of 200 μl was present in all wells except in blank where 100 μl media was added and the plates were incubated for 16 hours in an anaerobic workstation (A 35, Don Whitley, Yorkshire, UK). After the desired incubation period, the wells were washed thrice with phosphate buffer saline to remove the free-floating planktonic bacteria. The wells were subsequently stained with 100 μl crystal violet (0.4% w/v in distilled water) for 15 min. Following this, the wells were additionally washed thrice to remove any unbound crystal violet dye and were later dried for 2 h at 37 °C. Around 100μl of 95% (v/v) ethanol was added in each well to release the stain from the bacterial cells. The OD value was recorded at 630 nm using an ELISA plate reader (Multiskan™ FC Microplate Photometer, USA).

The absorbance in the blank well was subtracted from the absorbance reading of the test sample and the mean percent inhibition was calculated by:

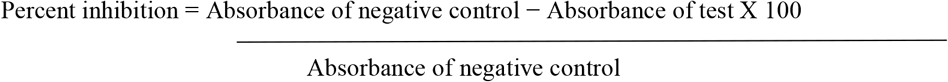

### 2.5 Anti-biofilm Assay

The anti-biofilm activity of goji berry extract was studied by adding 100 μl of Pg culture to each well in a 96-well microtitre plate and was incubated for 8 hours in an anaerobic workstation for the formation of biofilm. After 8 hours of incubation, 100 μl of test samples corresponding to their MBC values (Goji, CHX, and blank media) were added in triplicate to a 96-well microplate to make up the volume up to 200 μl. The microtitre plate was incubated for 48 hours in the anaerobic workstation. Crystal violet assay was carried out as previously mentioned and the inhibition of biofilm formation was assessed by the above-mentioned formula.

## 3. Results

### 3.1. Minimum Inhibitory Concentration

The LBE inhibited the growth of all the microorganisms as compared to the control (Table 1). The MIC was comparable to the gold standard (chlorhexidine) but higher as compared to the antibiotic doxycycline. The growth of one of the important periodontal pathogen (Pi) was reduced at a concentration of 3.12μg/ml. Pg and Aa required a greater concentration (6.25 μg/ml) to inhibit its growth. Fn and Tf were least sensitive to the extract and required a higher concentration of 50 μg/ml when compared to the other periodontal pathogens. At 50 μg/ml, all the periodontal pathogens were sensitive to the LBE compared to doxycycline (sensitive at 0.2 μg/ml), and chlorhexidine (sensitive at 32 μg/ml).

### 3.2 Minimum Bactericidal Concentration (MBC) (Table 1)

The LBE inhibited the colony formation for most of the pathogens (P.g, Aa, Pi) at 6.25 μg/ml with Tf and Fn requiring a higher concentration of 25 μg/ml. This value of MBC for LBE was comparable to chlorhexidine (8.3 μg/ml) and much higher when compared to doxycycline (0.2 μg/ml). Among the anaerobic bacteria, LBE seemed to be most effective for Pi.

### 3.3 Well Diffusion Assay (Figure 1 and Table 2)

The well-diffusion assay showed that LBE inhibited the growth of all the periodontal pathogen as compared to control. However, the zone of inhibition of the LBE was less when compared to doxycycline and chlorhexidine. The de novo extract showed a maximum zone of inhibition against Tf (15mm). Aa was found to be least inhibited by LBE.

### 3.4 Cell Proliferation assay (Figure 2)

LBE maintained the cell viability of more than 75 % of the gingival fibroblast and 80% of the modified keratinocyte cell line (HACAT) at a concentration of 1mg and 5 mg respectively compared to control.

### 3.5 Anti-adhesion Assay and Anti-biofilm Assay (Figure 3)

The results of the anti-adhesion showed that LBE (96%) was comparable compared to chlorhexidine (96.3%) However, it was noted that the anti-biofilm activity of chlorhexidine (96%) was found to be slightly better than that of Goji extract (91.6%).

## 4. Statistical analysis

The statistical analysis was done by the one-way ANOVA revealed an increase in the mean zones of inhibition against all the microorganisms which was statistically significant (P < 0.001). A comparison with doxycycline revealed insignificant inhibition zones by all the concentrations for all the pathogens. Post-hoc tests revealed a significant difference in the antimicrobial efficacy of doxycycline, chlorhexidine, and LBE against with mean zone of inhibition obtained by doxycycline and exceeding the mean zones of inhibition obtained by different concentrations of LBE and chlorhexidine. The zone of inhibition for chlorhexidine and LBE was comparable (p<0.001). The only exception LBE was found to be less effective /resistant to Aa as compared to chlorhexidine and doxycycline

## 5. Discussion

The results of the present study confirm that goji berry fruit extract has a good antimicrobial effect against oral periodontal pathogens. The extract also maintains the viability with gingival fibroblast and keratinocytes The MIC/MBC/ well diffusion assay should that LBE is sensitive to Pg, Tf, Aa, Fn, Pi, Sm. The MIC/MBC was comparable to chlorhexidine and much higher when compared to doxycycline. The MBC values for all microorganisms indicated that both chlorhexidine or LBE inhibited the growth of biofilm equally. Doxycycline was found to be superior to both LBE and chlorhexidine in inhibiting the growth of all the tested microorganisms. The results of the anti-adhesion and anti-biofilm assay also confirmed that the antiplaque efficacy of LBE was comparable to chlorhexidine. Therefore, LBE could serve as an alternative to Chlorhexidine for the management of periodontal diseases. The use of the herbal extract of goji berry instead of chlorhexidine will preclude adverse effects like tongue discoloration, tooth staining, and altered taste sensation. However, further in vivo studies and randomized clinical trials are required to confirm the efficacy of LBE as an adjunct to mechanical debridement for the management of chronic periodontitis.

These results of the antimicrobial, anti-adhesion and antibiofilm properties of LBE are attributed to its key constituents, which have strong antioxidant, antibacterial and anti-inflammatory properties (Kwok et al., 2019). Kabir et al. (2014) found that the chlorogenic acid in LBE can also contribute to its antimicrobial effects as it significantly increases the permeability of the cell and plasma membrane that can cause loss of the barrier function and leakage of cellular nucleotide that can cause cell death. The chlorogenic acid is confirmed to have both bacteriostatic and bactericidal effects against Gram-negative bacteria (*E. coli, Salmonella Typhimurium, Shigella dysenteriae*) and Gram-positive (*S. aureus, B. subtilis, S. pneumoniae*) (Li et al. 2006). Chlorogenic acid can also inhibit fatty acid synthase (FAS I) and the b-ketoacyl-ACP reductase (FabG) in a concentration-dependent manner, thereby reducing the pathogenic activity of *E. coli*. The rutin in the LBE has strong antibacterial activities as it can inhibit DNA isomerase IV (Orhan et al. 2010; Bernard et al. 1997). LBE also has an immunomodulatory effect, where it affects the cells of both innate and humoral immune response. An in-vitro mechanistic study conducted by Du et al. (2014) demonstrated that goji berry supplementation can enhance the maturation and activity of immune cells and change the production of inflammatory mediators. An upregulation in the recruitment of neutrophils and monocytes to sites of infection and an increase in phagocytosis has also been observed by LBE (Ren et al. 2012). These properties of LBP would be beneficial in lowering the microbial load and controlling the periodontal inflammation.

Based on these results and existing evidence, *Lycium barbarum* can be tried as a promising herbal alternative to manage oral and periodontal diseases. However, our study has evaluated the anti-microbial effects against the common periodontal pathogens, hence it is important to further explore the effects of LBE on other primary and secondary colonizers. It is also necessary to evaluate the effects of the individual constituents on oral microorganisms. *Lycium barbarum* can be incorporated into various pharmaceutics formulations like toothpaste, gels, mouthwash for the management of oral and periodontal infections and tested via various clinical trials to confirm its efficacy in the management of gingival and periodontal diseases.

## 6. Conclusion

*Lycium barbarum* is a promising herb for the management of oral and periodontal infections. The extract of *Lycium barbarum* is compatible with gingival fibroblast tissues and keratinocytes. The antimicrobial, anti-adhesion and antiplaque efficacy are comparable to chlorhexidine but less than that of doxycycline. *Lycium barbarum* could be used as a promising alternative to Chlorhexidine for the management of gingival and periodontal inflammation. However, further research and comparative clinical trials should be undertaken to evaluate and compare the effects of *Lycium barbarum* as an adjunct to mechanical debridement and a suitable alternative to a conventional antimicrobial agent for the management of periodontal diseases.

## 7. Acknowledgments

We would like to thank Dr. Kishore Bhat and his team members at Maratha Mandal’s Central Research Laboratory Maratha Mandal’s NGH Institute of Dental Sciences and Research Centre, Belgaum, for the antimicrobial analysis.

## References

1. Amagase, H., & Farnsworth, N. R., 2011. A review of botanical characteristics, phytochemistry, clinical relevance in efficacy, and safety of Lycium barbarum fruit (Goji). Food Res. Int. 44(7), 1702–1717. https://doi.org/10.1016/j.foodres.2011.03.027

2. Bautista-Pérez, R., Segura-Cobos, D., Vázquez-Cruz, B., 2004. In vitro antibradykinin activity of Aloe barbadensis gel. J Ethnopharmacol. 93(1), 89–92. https://doi.org/10.1016/j.jep.2004.03.030

3. Bell BG, Schellevis F, Stobberingh E, Goossens H, Pringle M. A systematic review and meta-analysis of the effects of antibiotic consumption on antibiotic resistance. BMC Infect Dis. 2014 Jan 9;14:13. doi: 10.1186/1471-2334-14-13. PMID: 24405683; PMCID: PMC3897982.

4. Belting, C.M., Massler, M., Schour, I., 1953. Prevalence and incidence of alveolar bone disease in men. J Am Dent Assoc, 47(2), 190–197. https://doi.org/14219/jada.archive.1953.0152

5. Bernard, F.X., Sable, S., Cameron, B., Provost, J., Desnottes, J.F., Crouzet, J., Blanche, F., 1997. Glycosylated flavones as selective inhibitors of topoisomerase IV. Antimicrob Agents Chemother. 41(5), 992–8. https://doi.org/10.1128/AAC.41.5.992

6. Bhat, G., Kudva, P., Dodwad, V., 2011. Aloe vera: Nature’s soothing healer to periodontal disease. J Indian Soc Periodontol. 15(3), 205–209. https://doi.org/10.4103/0972-124X.85661

7. Bondia-Pons, I., Savolainen, O., Törrönen, R., Martinez, J.A., Poutanen, K., Hanhineva, K., 2014. Metabolic profiling of Goji berry extracts for discrimination of geographical origin by non-targeted liquid chromatography coupled to quadrupole time-of-flight mass spectrometry. Food Res. Int. 63, 132–138. https://doi.org/10.1016/j.foodres.2014.01.067

8. Bonesvoll, P., Gjermo, P., 1978. A comparison between chlorhexidine and some quaternary ammonium compounds with regard to retention, salivary concentration and plaque-inhibiting effect in the human mouth after mouth rinses. Arch. Oral Biol. 23(4), 289–294. https://doi.org/10.1016/0003-9969(78)90021-3

9. Bonito AJ, Lux L, Lohr KN. Impact of local adjuncts to scaling and root planing in periodontal disease therapy: a systematic review. J Periodontol. 2005 Aug;76(8):1227-36. doi: 10.1902/jop.2005.76.8.1227. Erratum in: J Periodontol. 2006 Feb;77(2):326. Erratum in: J Periodontol. 2006 Feb;77(2):326–327.

10. Carrizales-Sepúlveda EF, Ordaz-Farías A, Vera-Pineda R, et al. Periodontal Disease, Systemic Inflammation and the Risk of Cardiovascular Disease. Heart Lung Circ 2018;27:1327–34.

11. Chapple ILC, Mealey BL, Van Dyke TE, et al. Periodontal health and gingival diseases and conditions on an intact and a reduced periodontium: Consensus report of workgroup 1 of the 2017 World Workshop on the Classification of Periodontal and Peri-Implant Diseases and Conditions. J Periodontol 2018;89 Suppl 1:S74–84.

12. Chen, Z., Soo, M.Y., Srinivasan, N., Tan, B.K.H., Chan, S.H., 2009. Activation of Macrophages by Polysaccharide–protein Complex from Lycium barbarum L. Phytother Res. 23(8),1116–22. https://doi.org/10.1002/ptr.2757.

13. Donno, D., Beccaro, G.L., Mellano, M.G., Cerutti, A.K., Bounous, G., 2015. Goji berry fruit (Lycium spp.): antioxidant compound fingerprint and bioactivity evaluation. J Funct Foods. 18, 1070–1085. https://doi.org/10.1016/j.jff.2014.05.020

14. Du, X., Wang, J., Niu, X., Smith, D., Wu, D., Meydani, S.N., 2014. Dietary wolfberry supplementation enhances the protective effect of flu vaccine against influenza challenge in aged mice. J Nutr. 144(2), 224–9. https://doi.org10.3945/jn.113.183566.

15. Flint, H. L. 1997. Landscape plants for eastern North America: exclusive of Florida and the immediate Gulf Coast. John Wiley & Sons.

16. Forino, M., Tartaglione, L., Dell’Aversano, C., Ciminiello, P., 2016. NMR-based identification of the phenolic profile of fruits of Lycium barbarum (goji berries). Isolation and structural determination of a novel N-feruloyl tyramine dimer as the most abundant antioxidant polyphenol of goji berries. Food Chem. 194, 1254–1259. https://doi.org/10.1016/j.foodchem.2015.08.129

17. Graves, D. 2008. Cytokines that promote periodontal tissue destruction. J Periodontol, 79(8), 1585–91. https://doi.org/10.1902/jop.2008.080183.

18. Kabir, F., Katayama, S., Tanji, N., Nakamura, S., 2014. Antimicrobial effects of chlorogenic acid and related compounds. J Korean Soc Appl Biol Chem 57(3), 359–365 (2014). https://doi.org/10.1007/s13765-014-4056-6

19. Karygianni L, Al-Ahmad A, Argyropoulou A, Hellwig E, Anderson AC, Skaltsounis AL. Natural Antimicrobials and Oral Microorganisms: A Systematic Review on Herbal Interventions for the Eradication of Multispecies Oral Biofilms. Front Microbiol. 2016 Jan 14;6:1529. doi: 10.3389/fmicb.2015.01529.

20. Keestra JA, Grosjean I, Coucke W, Quirynen M, Teughels W. Non-surgical periodontal therapy with systemic antibiotics in patients with untreated chronic periodontitis: a systematic review and meta-analysis. J Periodontal Res. 2015 Jun;50(3):294–314. doi: 10.1111/jre.12221.

21. Kumar, G., Jalaluddin, M.D., Rout, P., Mohanty, R., Dileep, C.L., 2013. Emerging trends of herbal care in dentistry. J Clin Diagn Res. 7(8), 1827–1829. https://doi.org/10.7860/JCDR/2013/6339.3282

22. Kushiyama, M., Shimazaki, Y., Murakami, M., Yamashita, Y., 2009. Relationship between intake of green tea and periodontal disease. J Periodontol. 80(3), 372–7. https://doi.org10.1902/jop.2009.080510.

23. Kwok, S.S., Bu, Y., Lo, A.C.Y., Chan, T.C.Y., So, K.F., Lai, J.S.M., 2019. A systematic review of potential therapeutic use of Lycium barbarum polysaccharides in disease. Biomed Res Int. https://doi.org/10.1155/2019/4615745

24. La, V.D., Howell, A.B., Grenier, D., 2010. Anti-Porphyromonas gingivalis and anti-inflammatory activities of A-type cranberry proanthocyanidins. J Antimicrob Chemother. 69(2), 428–436. https://doi.org/10.1093/jac/dkt398

25. Li, B.H., Ma, X.F., Wu, X.D., Tian, W.X., 2006. Inhibitory activity of chlorogenic acid on enzymes involved in the fatty acid synthesis in animals and bacteria. IUBMB life, 58(1), 39–46. https://doi.org/10.1080/15216540500507408

26. Low, K. 2019. Asian Fruits and Berries: Growing Them, Eating Them, Appreciating Their Lore. McFarland.

27. Mocan, A., Zengin, G., Simirgiotis, M., Schafberg, M., Mollica, A., Vodnar, D.C., Crişan, G., Rohn, S., 2017. Functional constituents of wild and cultivated Goji (L. barbarum L.) leaves: phytochemical characterization, biological profile, and computational studies. J Enzyme Inhib Med Chem. 32(1), 153–168. https://doi.org/10.1080/14756366.2016.

28. Nazir M, Al-Ansari A, Al-Khalifa K, Alhareky M, Gaffar B, Almas K. Global Prevalence of Periodontal Disease and Lack of Its Surveillance. Scientific World Journal. 2020;2020:2146160. doi:10.1155/2020/2146160

29. Orhan, D.D., Özçelik, B., Özgen, S. and Ergun, F., 2010. Antibacterial, antifungal, and antiviral activities of some flavonoids. Microbiol Res. 165(6), 496–504. https://doi.org/10.1016/j.micres.2009.09.002.

30. Pires, T.C., Dias, M.I., Barros, L., Calhelha, R.C., Alves, M.J., Santos-Buelga, C., Ferreira, I.C., 2018. Phenolic compounds profile, nutritional compounds and bioactive properties of Lycium barbarum L.: A comparative study with stems and fruits Ind Crops Prod. 122, 574–581. https://doi.org/10.1016/j.indcrop.2018.06.046

31. Ren, Z., Na, L., Xu, Y., Rozati, M., Wang, J., Xu, J., Sun, C., Vidal, K., Wu, D., Meydani, S.N., 2012. Dietary supplementation with lacto-wolfberry enhances the immune response and reduces pathogenesis to influenza infection in mice. J Nutr. 142(8), 1596–602. https://doi.org10.3945/jn.112.159467.

32. Skenderidis, P., Mitsagga, C., Giavasis, I., Petrotos, K., Lampakis, D., Leontopoulos, S., Hadjichristodoulou, C., Tsakalof, A., 2019. The in vitro antimicrobial activity assessment of ultrasound assisted Lycium barbarum fruit extracts and pomegranate fruit peels. J. Food Meas. Charact. 13(3), 2017–2031. https://doi.org/10.1007/s11694-019-00123-6.

33. Smiley CJ, Tracy SL, Abt E, Michalowicz BS, John MT, Gunsolley J, Cobb CM, Rossmann J, Harrel SK, Forrest JL, Hujoel PP, Noraian KW, Greenwell H, Frantsve-Hawley J, Estrich C, Hanson N. Systematic review and meta-analysis on the nonsurgical treatment of chronic periodontitis by means of scaling and root planing with or without adjuncts. J Am Dent Assoc. 2015 Jul;146(7):508–24.e5. doi: 10.1016/j.adaj.2015.01.028.

34. Toyoda-Ono, Y., Maeda, M., Nakao, M., Yoshimura, M., Sugiura-Tomimori, N., Fukami, H., 2004. 2-O-(β-D-Glucopyranosyl) ascorbic acid, a novel ascorbic acid analogue isolated from Lycium fruit J. Agric. Food Chem. 52(7), 2092–2096. https://doi.org/10.1021/jf035445w

35. Winning, L., Linden, G. Periodontitis and systemic disease. BDJ Team 2, 15163 (2015). https://doi.org/10.1038/bdjteam.2015.163

36. Xiao, X., Ren, W., Zhang, N., Bing, T., Liu, X., Zhao, Z., Shangguan, D., 2019. Comparative Study of the Chemical Constituents and Bioactivities of the Extracts from Fruits, Leaves and Root Barks of Lycium barbarum. Molecules, 24(8), 1585. https://doi.org/10.3390/molecules24081585

37. Zandbergen D, Slot DE, Cobb CM, Van der Weijden FA. The clinical effect of scaling and root planing and the concomitant administration of systemic amoxicillin and metronidazole: a systematic review. J Periodontol. 2013 Mar;84(3):332–51. doi: 10.1902/jop.2012.120040.

38. Zhang, Q., Chen, W., Zhao, J., Xi, W., 2016. Functional constituents and antioxidant activities of eight Chinese native goji genotypes. Food Chem. 200:230–236. https://doi.org/10.1016/j.foodchem.2016.01.046.

